# *De novo* design of phosphotyrosine peptide binders

**DOI:** 10.1101/2025.09.29.678898

**Authors:** Magnus S Bauer, Jason Z Zhang, Kejia Wu, Gyu Rie Lee, Brian Coventry, Isabella M Silvestri, Kody A Klupt, Jiuhan Shi, Rafael I Brent, Xinting Li, Carolina Moller, Nicole Roullier, Dionne K Vafeados, Indrek Kalvet, Rebecca K Skotheim, Siyu Zhu, Amir Motmaen, Luca C Herrmann, Pascal Sturmfels, Doug Tischer, Han Altae-Tran, David Juergens, Rohith Krishna, Woody Ahern, Jason Yim, Asim K Bera, Alex Kang, Emily Joyce, Andrew Lu, Lance Stewart, Frank DiMaio, Krishna C Mudumbi, David Baker

**Affiliations:** Department of Biochemistry, University of Washington, Seattle, WA 98195, USA; Institute for Protein Design, University of Washington, Seattle, WA 98195, USA; Department of Bioengineering, University of California Los Angeles, Los Angeles, CA 90095, USA; Department of Biological Sciences, Korea Advanced Institute of Science and Technology, Daejeon, South Korea; Department of Cell and Developmental Biology, Vanderbilt University, Nashville, TN 37232, USA; The Vanderbilt-Ingram Cancer Center, Vanderbilt University Medical Center, Nashville, TN 37232, USA; Department of Biomedical and Molecular Sciences, Queen’s University, Kingston, ON, Canada; Computer Science and Artificial Intelligence Laboratory, Massachusetts Institute of Technology, Cambridge, MA 02139, USA; Department of Biology and Biological Engineering, California Institute of Technology, Pasadena, CA 91125, USA; Howard Hughes Medical Institute, University of Washington, Seattle, WA 98195, USA

## Abstract

Phosphorylation on tyrosine is a key step in many signaling pathways. Despite recent progress in *de novo* design of protein binders, there are no current methods for designing binders that recognize phosphorylated proteins and peptides; this is a challenging problem as phosphate groups are highly charged, and phosphorylation often occurs within unstructured regions. Here we introduce RoseTTAFold Diffusion 2 for Molecular Interfaces (RFD2-MI), a deep generative framework for the design of binders for protein, ligand, and covalently modified protein targets. We demonstrate the power and versatility of this method by designing binders for four critical phosphotyrosine sites on three clinically relevant targets: Cluster of Differentiation 3 (CD3ε), Epidermal Growth Factor Receptor (EGFR), Insulin Receptor (INSR) and Signal Transducer and Activator of Transcription 5 (STAT5). Experimental characterization shows that the designs bind their phosphotyrosine containing targets with affinities comparable to native binding sites and have negligible binding to non-phosphorylated targets or phosphopeptides with different sequences. X-ray crystal structures of generated binders to CD3ε and EGFR are very close to the design models, demonstrating the accuracy of the design approach. A designed binder to an EGFR intracellular region phosphorylated upon EGF activation co-localizes with the receptor following EGF stimulation in single-particle tracking (SPT) experiments, demonstrating pY specific recognition in living cells. RFD2-MI provides a generalizable all-atom diffusion framework for probing and modulating phosphorylation-dependent signaling, and more generally, for developing research tools and targeted therapeutics against post-translationally modified proteins.

## INTRODUCTION

Phosphorylation is one of the most critical post-translational modifications (PTMs) in cell signaling and other cellular processes. To probe and modulate phosphorylation sites, phosphorylation-specific binding reagents are needed, yet current options are limited. Antibodies, the most commonly used class of binders, often fail to distinguish between closely related phosphosites (1–3), require long development times, and are constrained by a single scaffold architecture that limits the diversity of interaction surfaces. Directed evolution strategies using nature-derived domains that bind to phosphosites like Src Homology 2 (SH2) or WW domains can improve general phospho-residue binding (e.g. phosphotyrosines (pY), phosphothreonines or phosphoserines), but lack the ability to discriminate between sequences flanking the phosphorylation site (4–7). This limitation is especially problematic for disordered protein regions, where phosphorylation often occurs, but where conventional binding domains have difficulty achieving specificity due to the absence of structured binding pockets. These challenges underscore the need for a fundamentally new approach to develop small, robust, and sequence-specific phospho-binders that can function inside living cells and serve as precision tools for sensing or modulating phosphorylation dynamics.

Deep learning based design of proteins has revolutionized the ability to create binders to challenging protein targets, but general methods for designing binders to covalently modified proteins and peptides have not been described. We reasoned that RoseTTAFold Diffusion2 (RFD2), a recently developed method for enzyme and small molecule binder design, could be generalized to designing binders for phosphotyrosine-containing (pY) peptides by training jointly on complexes of proteins with other proteins, small molecules, and with covalently modified proteins and peptides. We set out to extend RFD2 for binder design against this full range of targets and to explore the use of the new method for designing binders to pY containing peptides.

## RESULTS

### Designing binders to covalently modified proteins using RoseTTAFold Diffusion 2 for molecular interfaces (RFD2-MI)

RFD2 extended the backbone-level RFdiffusion framework to enable generation of proteins making specific sidechain-small molecule interactions using a flow-matching denoising objective (8, 9). RFD2 can design proteins with precise small-molecule binding sites for enzymes, using unindexed and tip-atom constraints to position catalytic groups at full atomic resolution without pre-specifying sequence placement. These capabilities make it highly effective for active-site scaffolding, but they are tailored to enzyme-ligand contexts. To enable binder design for protein-protein interactions or protein-ligand interfaces with defined binding sites and steerable interface properties we trained a new model based on RFD2 called RFD2 for molecular interfaces (RFD2-MI). We include additional interface examples and inject specialized 1D features that allow us to conditionally modify both the binder structure and the corresponding binding site on the target. The 1D feature tensor in RFD2-MI is provided as a per token embedding (tokens are residues or atoms) that can encode amino-acid identity (one-hot), secondary-structure tokens (helix/strand/coil), motif flags, hotspot/antihotspot masks, solvent-accessibility, and any user-supplied conditions. At each diffusion step, RFD2-MI concatenates a per-residue 1D feature vector and then broadcasts that embedding to all of the residue’s atom slots, so every coordinate update is conditioned on those features (**Figure 1A**).

**Figure 1.**
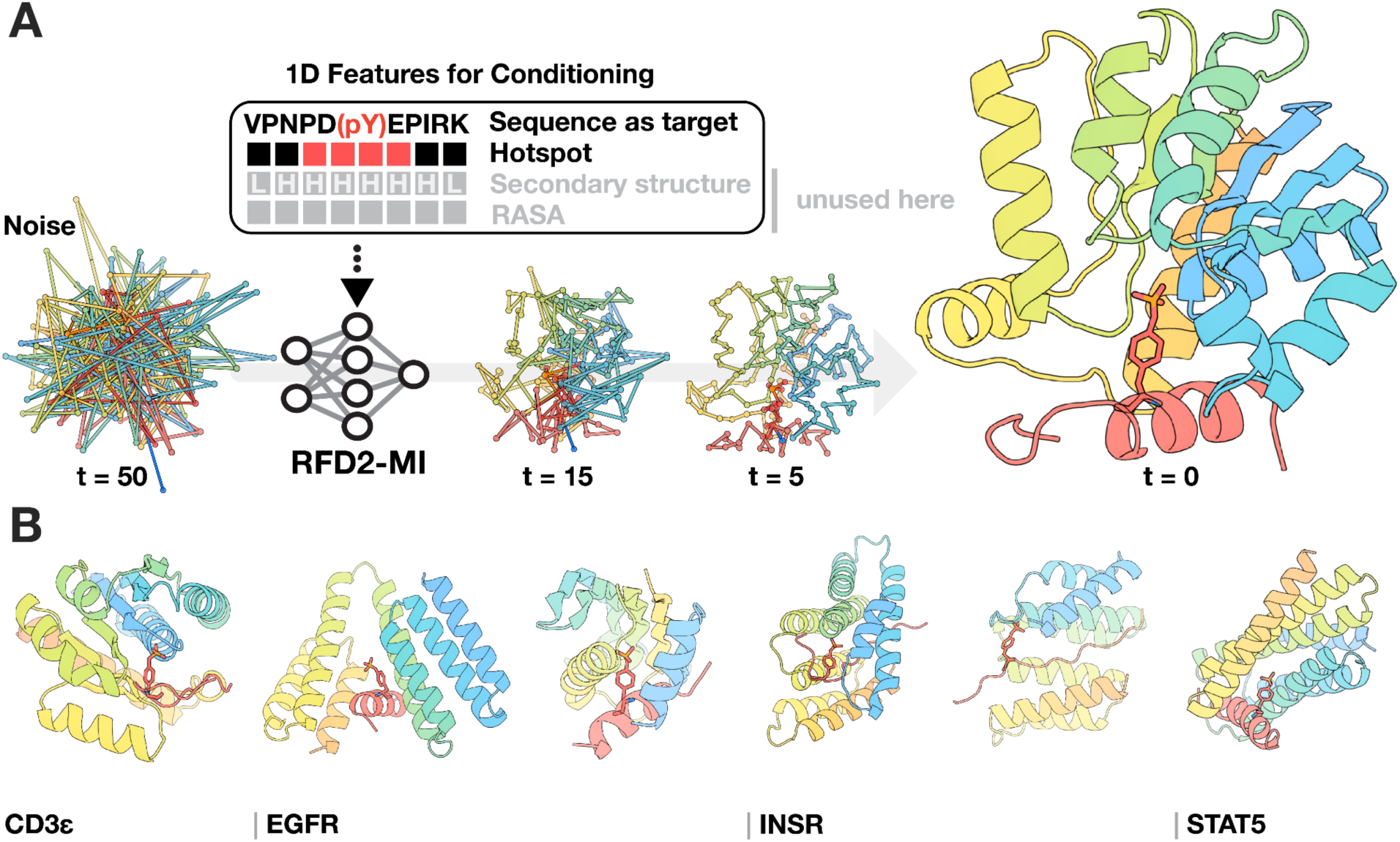
RFD2-MI enables *de novo* design of phosphotyrosine-specific peptide binders from target sequence alone. **(A)** Schematic of the RFD2-MI workflow. **(top)** The 1D conditioning feature channels can include hotspot/anti-hotspot masks, setting secondary structure, and solvent accessibility (RASA); in the examples shown here we design for a given peptide sequence, specified by sequence alone and conditioning on a hotspot mask (grayed items unused). Conditioning steers the model to form a pocket that positions the phosphotyrosine (pY; red/orange sticks in the peptide) and shapes complementary contacts in the designed binder (rainbow cartoon). **(bottom)** At each denoising step, the coordinates are diffused from noise (timesteps shown: t = 50, 15, 5, 0) while the binding site is guided through 1D conditioning features that are concatenated to the residue/atom embeddings. **(B)** Representative design outcomes for the three targets studied in this paper. RFD2-MI generated compact binders that cradle the cognate pY-containing peptide for CD3ε, EGFR, INSR (coupled with the *Logos* pipeline) and STAT5. In all cases the phosphorylated residue (red/orange, shown as sticks) is coordinated by a pocket in the *de novo* binder (rainbow), while neighboring sequence-specific contacts help encode selectivity.

RFD2-MI was trained on a curated, interface-focused dataset enriched for compact, well-packed binders in contact with their targets, while discarding poorly folded examples and trimming disordered or non-interacting regions for the binder (peptide targets are not trimmed). Hotspot and anti-hotspot masks guide interface geometry and prioritize key contacts. Binder placement is steered by the origin (ORI) of the starting residue Gaussian noise cloud, with optimal positioning achieved by centering it on the hotspot mask or near the intended binding site, yielding robust and shape-complementary designs (see Methods for details on training).

### Computational design and characterization of phosphorylation-specific binders

We used RFD2-MI (**Figure 1A, B**) to design binders to phosphopeptide targets central to cellular signaling: 1) CD3ε, a component of the T-cell receptor, 2) two sites on epidermal growth factor receptor (EGFR), a potent oncogene, and 3) insulin receptor (INSR), a regulator of glucose metabolism dysregulated in diabetes. Because these phosphorylations take place within unstructured regions, we co-diffuse the designed protein and the target peptide, allowing the peptide to explore conformations that align with complementary binding pockets in the designed protein. In the case of INSR, we coupled this pipeline with the previously published Logos (12) method to integrate an interface for the disordered substrate (see Methods). Following structure generation, LigandMPNN was used to optimize the sequence for high affinity binding to the modified peptides. Designs were filtered based on the change in Gibbs free energy (ΔΔG) computed using Rosetta (10) for overall binding, the number of hydrogen bonds between the side chains of target and designed protein (for sequence specificity), and the number of hydrogen bonds between designed protein and phosphate (for phosphorylation specificity). For the INSR binders, we utilized the Logos pipeline (11, 12) to generate binders for adjacent non-modified regions in the target, and then used RFD2-MI to extend these to bind the full target including the pY modified region (see Methods for details). Designs for which AF3 predictions of the binder-target complex were close to the generated backbones after Rosetta FastRelax and for which the sidechain phosphate hydrogen bonds were recapitulated were selected for experimental validation.

We obtained synthetic DNA encoding 459 designs for CD3ε pY188, 5,082 designs for EGFR pY1068, and 2,408 designs for EGFR pY1173, displayed the proteins on yeast, identified binders to the phosphorylated target peptides using yeast display (see Methods), and measured the binding affinities of *E coli* expressed protein to target peptide using biolayer interferometry (BLI). The designed CD3ε pY188 binder (CD3ε_pY188_bp) had a Kd of 2.3 μM for the phosphorylated CD3ε pY188 peptide target and undetectable binding for the non-phosphorylated version of the peptide (**Figure 2A-C, row 1**). The designed EGFR pY1068 binder (EGFR_pY1068_bp) had a Kd of 4.2 µM for the phosphorylated target and much weaker affinity for the nonphosphorylated peptide (**Figure 2A-C, row 2**). The designed EGFR pY1173 binder (EGFR _pY1173_bp) bound preferentially to the phosphorylated peptide with a Kd of 659 nM (**Figure 2A-C, row 3, S1**). For the designed INSR pY1316 binder (INSR_pY1316_bp), we obtained genes encoding 80 designs and expressed these directly in *E coli*. 71/80 of these expressed solubly; 5/80 designs bound to both the phosphorylated and non-phosphorylated peptide, and 2/80 designs bound considerably more strongly to phosphorylated target peptide than the non-phosphorylated peptide (INSR_pY1316_bp binds target with a Kd of 1.2 μM; **Figure 2A-C, row 4, S1**). In the case of the STAT5 pY694 target we obtained synthetic DNA encoding 2,835 designs and identified 4 binders by yeast display. The lead designed, STAT5 pY694 binder (STAT5_pY694_bp), bound the phosphorylated STAT5 peptide with a Kd of 317 nM determined by BLI (**Figure 2A-C, row 5; Figure S1**). Lead binders show sub-µM to low-µM affinities, in the range reported for many endogenous phosphotyrosine-recognition interactions (13, 14).

**Figure 2.**
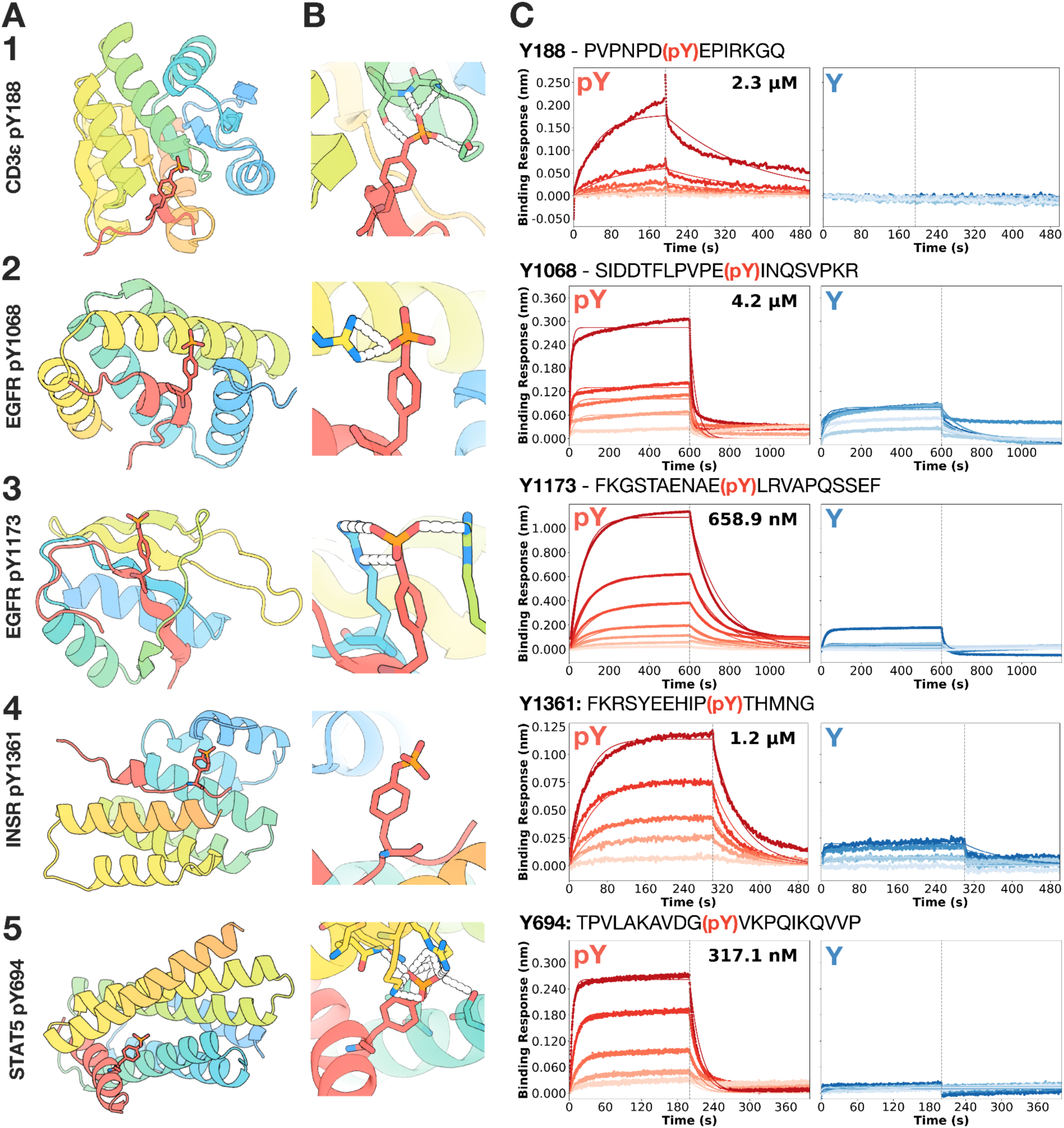
Structural and biophysical characterization of phosphopeptide-specific binders. **(A)** Design models (rainbow) in complex with the four target peptides: CD3ε pY188 (row 1), EGFR pY1068 (row 2), EGFR pY1173 (row 3), INSR pY1361 (row 4) and STAT5 pY694 (row 5). In each case the phosphorylated peptide (red/orange sticks) docks into the designed pocket. **(B)** Close-up views of key hydrogen-bonding networks (dashed lines) in the design model for CD3ε pY188, EGFR pY1068, EGFR pY1173, INSR pY1361 and STAT5 pY694 with their respective binders, highlighting how the phosphotyrosine moiety (red/orange sticks) is anchored by a conserved polar pocket in the binder (rainbow). **(C)** Representative biolayer interferometry (BLI) sensorgrams showing association and dissociation of each binder to phosphorylated (blue) versus non-phosphorylated (red) peptides, with global 1:1 kinetic fits overlaid in black. CD3ε_pY188_bp, EGFR_pY1068_bp, EGFR_pY1173_bp, INSR_pY1361_bp and STAT5_pY694_bp exhibit dose-dependent binding only to the phosphorylated peptide, showing a much more obvious binding toward pY peptides than their unmodified counterparts.

For CD3ε_pY188_bp and EGFR_pY1068_bp, the phosphopeptide adopts a partially ɑ-helical conformation with the phosphate at the end of the ɑ-helix and the phosphate interacts with a loop region in the binder. CD3ε_pY188_bp coordinates the target phosphate group with the backbone atoms of a p-loop at the terminus of a helix, similar but not identical to the Walker motif (15, 16) associated with phosphate binding and coordination of ATP. For EGFR_pY1173_bp, the phosphopeptide adopts a beta strand that pairs with a beta sheet in the binder; the phosphate is in the middle of the β-sheet, and the phosphate interacts with the side chains of residues in the neighboring β-sheets (**Figure 2A-C, row 3**). For the Logos-derived INSR designs, the target has an extended confirmation and makes extensive polar interactions with the binder particularly around the charged phosphate group (**Figure 2A-C, row 4**), which likely confers the bias towards the phosphorylated version of the target.

### Structural validation of phosphorylation-specific binders

The crystal structures of both the CD3ε pY188 and EGFR pY1068 binder-target complexes showed strong agreement with the design models (**Figure 3**). For the CD3ε pY188 complex, the overall complex Cα RMSD between the crystal structure and the design model is 2.1 Å, indicating the accuracy of the design methods. The phosphotyrosine was very accurately placed; following the superposition of the crystal structure and design model based on the binder, the RMSD on the phosphotyrosine was only 1.91 Å, demonstrating the capabilities of the model to design well coordinated phosphate binding motifs. For the EGFR pY1068 complex, structural agreement was also quite high, with a complex Cα RMSD of 2.2 Å relative to the design model. The phosphate in the model aligns well with the crystal structure, with an RMSD of 2.4 Å, indicating highly precise phosphorylated residue placement during design. The phosphate-binding pocket retained all intended hydrogen-bonding interactions, with the phosphate coordinated by two arginine side chains and the backbone amide of a nearby alanine residue. The relative orientation of the peptide and binder was essentially unchanged, and the local geometry around the phosphate group was reproduced with sub-angstrom accuracy. Taken together, these results highlight the accuracy of RFD2-MI-guided design in capturing not only the global fold of the binder-target complex but also the precise positioning of phosphate-binding motifs.

**Figure 3.**
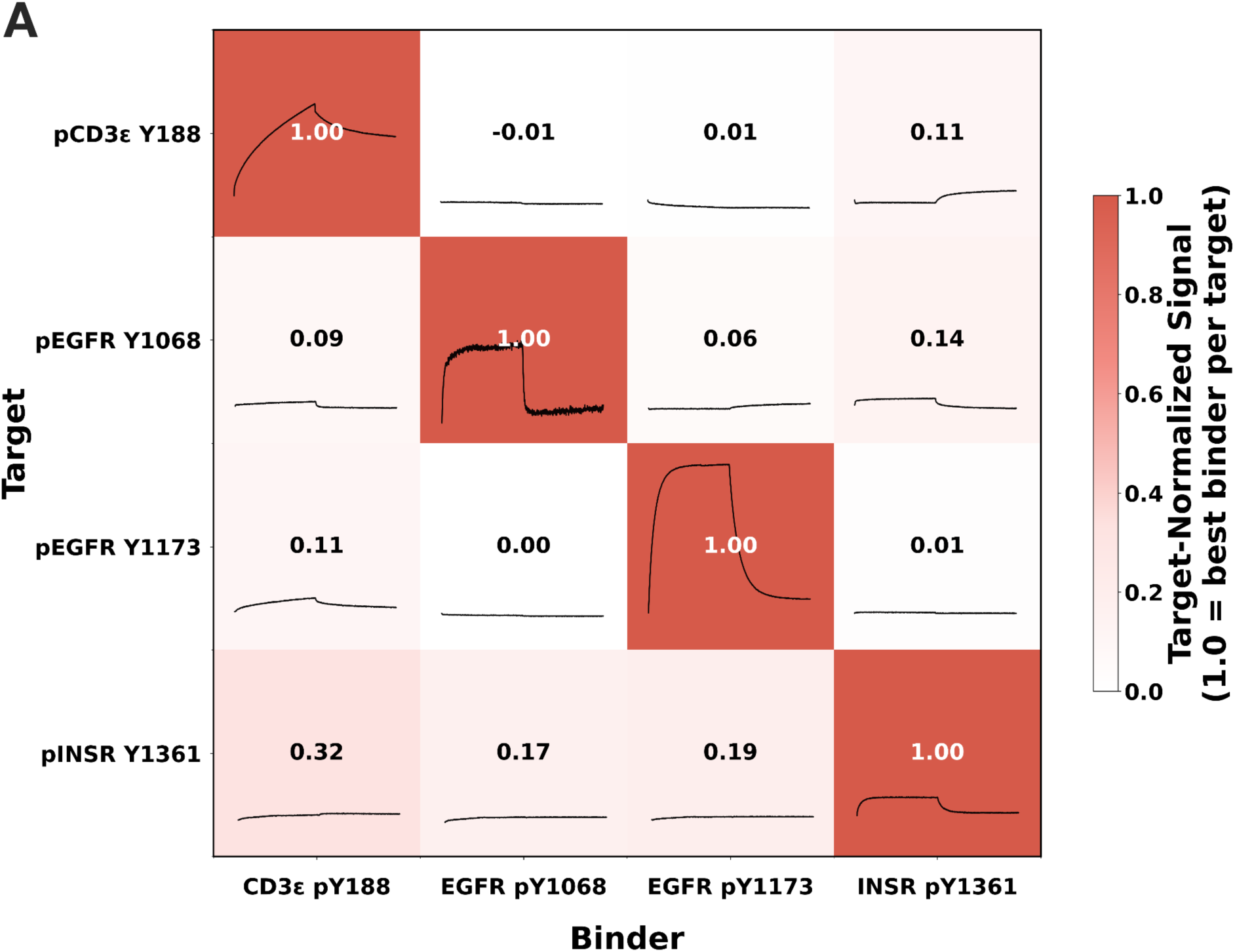
Crystal structures of phosphorylation-specific binders match the design models. **(A)** Overlays of the crystal structures (rainbow) with design models (gray) for CD3ε_pY188_bp (top) and EGFR_pY1068_bp (bottom). Black boxes indicate the binding pockets. The experimental structures closely follow the design models. **(B)** Zoom-in views of the phosphate-recognition motifs with electron density shown as a blue mesh (2Fo–Fc map contoured at 2.5σ). The phosphotyrosine (PTR) is displayed in sticks and contacting binder residues are labeled. For CD3ε_pY188_bp, a short p-loop-like segment (G63-A67) coordinates the phosphate, capped by R69. For EGFR_pY1068_bp, the phosphate is chelated by two arginines (R7, R83) and stabilized by the nearby backbone amide of A10, illustrating the intended polar pocket that underlies phosphorylation specificity.

### Evaluation of sequence specificity

With many phosphorylations occurring in a cell at any given time, differentiating between different phosphorylation sites is critical to understanding biological mechanisms. Ideally, our designed binders should be both phosphorylation and sequence specific to differentiate phosphorylation sites. We encoded sequence specificity in our design pipeline in several ways. Structural sampling of both the designed protein and peptide target during RFD2-MI enables the designed protein to create a binding pocket for a particular peptide conformation, thus maximizing interactions. In addition, filtering for designs with multiple hydrogen bonds between the side chains of the design and target boosts the probability for sequence specificity as side chain-side chain interactions are sequence specific. Indeed, the designed pY peptide binders make extensive polar interactions with their target, which we anticipated would lead to sequence specificity. We assessed sequence specificity using an all-by-all BLI assay (**Figure 4**), measuring binding of four designs against each of the four targets against all four phosphopeptides. Strong signals were observed only for the cognate design-peptide pairs, with negligible signal for non-cognate pairs. These results demonstrate that the designs have sequence specificity.

**Figure 4.**
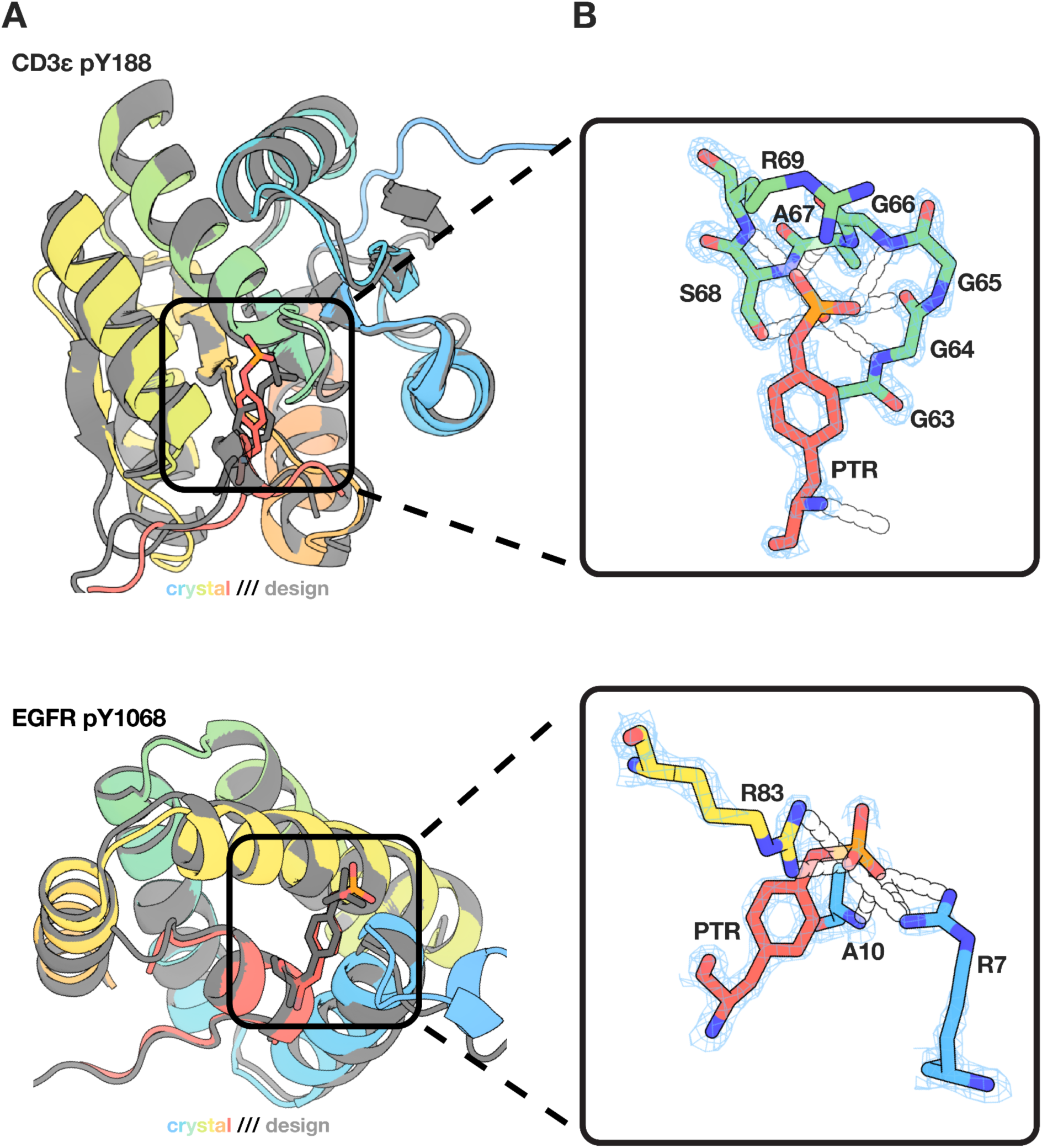
Designed binders have sequence specificity. **(A)** All-by-all binding heatmap displaying bio-layer interferometry (BLI) binding traces and normalized signal intensities. Phospho-binders (x-axis) were tested against target phosphopeptides (y-axis), with embedded kinetic traces showing association and dissociation profiles at the highest tested concentration. Color intensity and numerical values represent binding strength normalized to the strongest interaction per row (1.00 = 100% of maximum signal per target peptide). The matrix reveals specificity patterns, with strongest signals along the diagonal indicating preferential binding of each phospho-binder to its cognate target, while off-diagonal interactions demonstrate negligible cross-reactivity levels.

### EGFR_pY1068_bp is recruited to phosphorylated EGFR upon EGF stimulation in living cells

We used single-particle tracking (SPT) to study the recruitment of individual peptide binders to their intended target, focusing on EGFR pY1068. To visualize the binding of our *de novo* generated EGFR_pY1068_bp to activated full-length EGFR in living cells, we conjugated the phospho-binder to HaloTag7 (EGFR_pY1068_bp^HT7^) and co-expressed it alongside EGFR-GFP in Chinese hamster ovary (CHO) cells, which lack endogenous EGFR. Using total internal reflection fluorescence (TIRF) microscopy we imaged cells expressing these constructs with and without saturating levels of EGF (50 nM), which is known to phosphorylate EGFR at residue Y1068 (18) (**Figure 5A)**. We found that cells treated with ligand exhibited an increase in the number of EGFR_pY1068_bp^HT7^ particles observed at the plasma membrane (**Figure 5B and C**). As a negative control, we expressed only the EGFR_pY1068_bp^HT7^ construct in CHO cells without co-expressing EGFR. Under these conditions we observed no change in the number of particles at the membrane in cells treated with EGF, indicating the observed recruitment is due to both the presence and activation of EGFR (**Figure 5C**).

**Figure 5.**
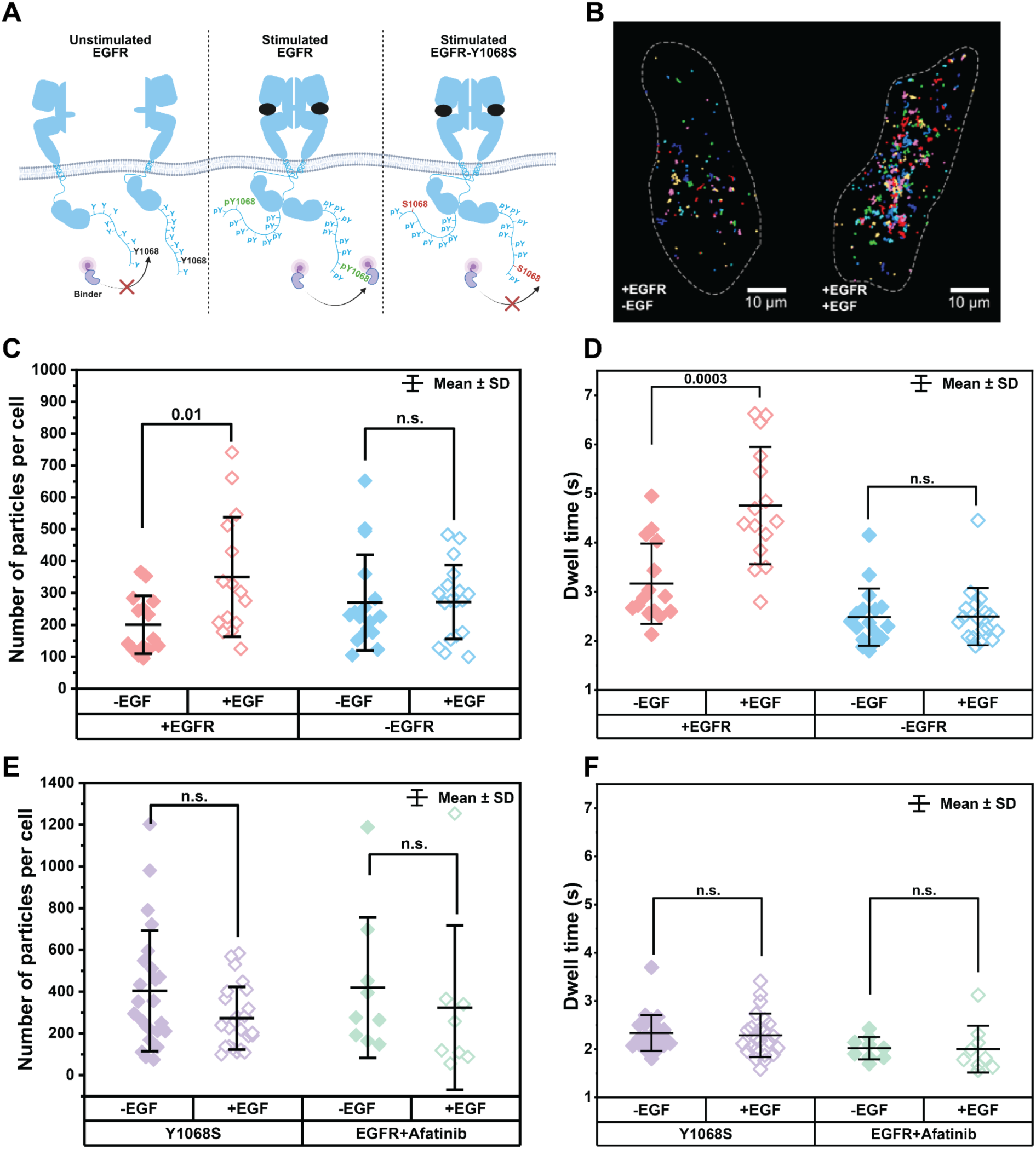
Phospho-binder EGFR_pY1068_bp is specific for phosphorylated Y1068 in living cells. **(A)** Schematic showing phospho-binder EGFR_pY1068_bp^HT7^ recruitment to EGFR in various conditions: unstimulated wild-type EGFR, stimulated wild-type EGFR, and stimulated EGFR^Y1068S^. **(B)** Representative images of EGFR_pY1068_bp^HT7^ recruitment to the cell surface in EGFR null CHO cells overexpressing wild-type EGFR either unstimulated or stimulated with 50 nM of EGF. **(C)** SPT of EGFR_pY1068_bp^HT7^ in cells expressing EGFR (pink), or not expressing EGFR (blue) when unstimulated (solid-diamonds) or stimulated with EGF (open-diamonds). Each point corresponds to the total number of tracks detected per cell per condition in SPT experiments. **(D)** Dwell times of tracked EGFR_pY1068_bp^HT7^ for conditions where it is co-expressed with wild-type EGFR (pink), or without co-expressing EGFR (blue) in CHO cells. Closed symbols represent cells without EGF stimulation, and open symbols represent cells stimulated with 50 nM of EGF. Each data point is the weighted average dwell time for all movement classes determined by DC-MSS analysis (17) of EGFR_pY1068_bp^HT7^ detected at the plasma membrane in each cell for each condition. EGFR_pY1068_bp^HT7^ molecules at the plasma membrane in the absence of ligand and in the absence of EGFR are primarily in the immobile fraction (**Figures S2 and S3**) and may reflect nonspecific interactions with the plasma membrane and/or other molecules at the plasma membrane. **(E)** SPT of EGFR_pY1068_bp^HT7^ in cells expressing EGFR with a mutation at tyrosine 1068 to serine (purple), or cells expressing wild-type EGFR treated with 10 µM of the tyrosine kinase inhibitor, afatinib (aquamarine). Open symbols represent conditions where cells were not stimulated with EGF, and closed symbols represent cells stimulated with 50 nM EGF. **(F)** Dwell times of tracked EGFR_pY1068_bp^HT7^ in cells expressing the mutant EGFR^Y1068S^ (purple), or wild-type EGFR treated with afatinib (aquamarine). Open symbols represent unstimulated cells, and closed symbols represent cells treated with 50 nM EGF. Unpaired two-sided Welche’s t-tests were used to calculate p-values. Significant p-values are represented above the data and n.s. indicates not significant at p = 0.05.

Having observed that EGFR_pY1068_bp^HT7^ is recruited to the plasma membrane in response to stimulation by ligand we next sought to understand the duration of binder interactions at the membrane through the particle dwell time. We used Divide-and-Conquer Moment Scaling Spectrum transient diffusion analysis (DC-MSS) (17) to classify portions of trajectories as free diffusion, confined diffusion, and immobile diffusion (**Figure S2**). We then parsed out the dwell times of binders within each class using weighted averages to determine the contribution of each diffusion class. In cells co-expressing EGFR_pY1068_bp^HT7^ and EGFR, there was an increase in phospho-binder dwell times at the plasma membrane (4.76 ± 1.2 seconds) when cells were stimulated with EGF (**Figure 5D)**. In contrast, in CHO cells where EGFR was not co-expressed with the binder, there was no change in EGFR_pY1068_bp^HT7^ dwell times at the plasma membrane with or without EGF (**Figure 5D**).

To evaluate the specificity of EGFR_pY1068_bp for the phosphorylated tyrosine residue at 1068 on EGFR, we mutated tyrosine 1068 to a serine (EGFR^Y1068S^). We then assessed how this affected EGFR_pY1068_bp^HT7^ recruitment to the plasma membrane and the dwell time of recruited particles. We observed little change in binder recruitment to the plasma membrane following stimulation with EGF (**Figure 5E**), and there was no change in EGFR_pY1068_bp^HT7^ dwell time at the plasma membrane upon EGF stimulation. We further tested the effects of treating cells with 10 µM of afatinib, a covalent tyrosine kinase inhibitor specific for EGFR that blocks EGFR phosphorylation. We again observed little change in binder recruitment to the plasma membrane or EGFR_pY1068_bp^HT7^ dwell time at the plasma membrane (**Figure 5F)**. Together these results indicate that the *de novo* phosphopeptide binder to EGFR pY1068 is specific for the phosphorylated Y1068 residue on the C-terminus of EGFR in living cells: only when wild-type EGFR is stimulated with ligand is there is an increase in EGFR_pY1068_bp^HT7^ recruitment to the plasma membrane with a significantly increased dwell time. The increase in dwell time for EGFR_pY1068_bp^HT7^ when co-expressed with EGFR but not stimulated with EGF (3.12 ± 0.82 seconds; **Figure 5D, and S3**) over cells not expressing EGFR, expressing EGFR^Y1068S^, or expressing wild-type EGFR but treated with afatinib, is consistent with previous findings that overexpression of EGFR leads to increased background phosphorylation of the receptor even without ligand stimulation (18, 19).

## DISCUSSION

By introducing conditioning features in RFD2 geared towards designing molecular interactions and the implementation of PTM-aware modeling, we enabled the model to generate binders that precisely recognize phosphorylated targets. This enhanced model, called RFD2-MI, successfully generated compact and diverse binders to multiple phosphopeptides. The two crystal structures of designs in complex with their targets illustrate the accuracy of the design approach, which is notable given the challenges in binding the highly charged phosphate group. In contrast to previous affinity reagents such as antibodies, our *de novo* designs are not confined to a particular structural scaffold (12, 20); although RFD2-MI was trained on the PDB, the current model does not simply recapitulate designs with phospho-binding motifs like SH2 or 14-3-3 domains. This may be particularly useful when designing specific binders against phosphopeptide targets that do not have sequence similarity with phosphopeptides bound by natural proteins. More generally, the ability of RFD2-MI to generate binders to proteins, small molecules, and covalently modified proteins with state of the art performance makes it applicable to a very broad class of binder design problems.

While RFD2-MI has demonstrated success in designing binders to phosphotyrosines, there is room for improvement in our computational design pipeline as the success rate is low (often <0.1%) and the affinities of the best binders are modest (above 500nM). Overall, the challenge of achieving specificity for the phosphorylated versus unphosphorylated peptide was greater than the challenge of achieving sequence specificity: we obtained many designs that bound both phosphorylated and non-phosphorylated forms, while all phosphorylation-specific binders were also sequence-specific. This is not surprising given the challenge in binding the highly charged and solvated phosphate group. To achieve increased affinity with the phosphate present requires more than compensating for the lost interactions with solvent upon binding. The relatively low affinities of the best binders likely derive in part from the cost of phosphate desolvation; the two crystal structures show that the designs are quite accurate structurally. Future phosphopeptide binder design could benefit from greater focus on optimizing the interactions with the phosphate, while reducing slightly the emphasis on neighboring sequence contacts. More generally, incorporation of multi-modal training data such as chemical, structural, and dynamic information (21) could further improve binder selectivity and functionality.

Our SPT experiments demonstrate that EGFR_pY1068_bp is recruited to the phosphorylated tyrosine residue 1068 on EGFR in response to stimulation by ligand. This demonstrates the utility of these designs to study previously difficult and/or inaccessible questions regarding receptor activation and signaling kinetics; for example, studying EGFR phosphorylation lifetime following stimulation by different ligands. Our binder dwell time studies suggest that Y1068 on EGFR is phosphorylated for about 4.5 seconds in response to EGF activation, providing higher temporal resolution than previous methods and studies (22–24). More generally, designed phosphopeptide binders should help elucidate new phenomena regarding the role of PTMs in cellular signaling.

Our RFD2-MI approach can be readily generalized to other PTMs such as methylation, acetylation, or glycosylation to enable the generation of custom binders for a wide range of regulatory protein states previously considered undruggable (25, 26). These binders could enable researchers to build precise tools for detecting and manipulating individual sub-steps within signaling cascades, such as ligand engagement, adaptor recruitment, kinase activation, or effector binding. By targeting these discrete events, it should become possible to map signaling dynamics with higher resolution and uncover previously hidden regulatory checkpoints; such capabilities would not only improve our ability to sense and perturb signaling in living cells but also open new strategies for therapeutic intervention. Ultimately, our approach could transform how protein modifications are studied and exploited across biology and medicine.

## METHODS

### All-atom molecular interaction model training

RFD2-MI extends RFD2 to all-atom generative modeling of molecular interfaces between proteins and ligands. RFD2-MI uses the SE(3) flow matching architecture established in RFD2, with Gaussian probability paths for translations and Riemannian flow for rotations, to iteratively denoise heavy-atom coordinates and adds per token 1-D features for adding conditioning to get molecular interactions of choice and additional cropping strategies for training examples.

RFD2-MI was trained with data sampling probabilities biased toward protein-protein complexes to 53% in order to enrich interface examples (27). Complex-like datasets were filtered to enforce a maximum binder length of 170 residues to control memory usage and match downstream cropping behavior. Folding quality was enforced via side-chain neighbor (SCN) analysis, where each binder residue’s packing density was measured using a virtual Cβ cone and Cα neighbors. Two folding metrics were computed: binder_FBSCN (fraction of all binder residues with SCN > 4.0) and interface_FBSCN (same fraction but restricted to binder interface residues). Examples were rejected if binder_FBSCN < 0.12, interface_FBSCN < 0.13, or no hotspots were detected. To further improve physical plausibility, disordered N/C-terminal binder tails were trimmed using a 9-residue sliding window tolerance scheme, allowing removal of floppy segments even into hotspot regions if necessary. These steps, combined with a radial crop around hotspots, produced a dataset biased toward compact, interface-packed binders with well-structured interfaces, while excluding unfolded or non-contacting complexes for the binder. The targets are not assessed for packing and also not trimmed so peptide targets will stay intact for training purposes.

To design binders against a target surface, we diffused only the binder while keeping the target fixed, conditioning on interface cues derived from hotspots and antihotspots on the target. A target residue was labeled as a hotspot if its Cβ lay within 7 Å of any binder Cβ and its solvent-accessible surface area was at least 30 Å² (probe radius 2.8 Å). Hotspots were present in 75% of training examples and downsampled to at most 20% coverage using a speckle-or-region masking strategy, with optional floating-point “hotspot values” representing the local density of neighbors within 10 Å to emphasize high-contact “super-hotspots.” Antihotspots were target residues more than 10 Å from any binder residue and were included in 10% of examples to discourage spurious contacts. Super hotspots augment the binary hotspot mask with a continuous weight per residue that reflects local contact density: for each hotspot that passes the 7 Å Cβ-Cβ proximity and 30 Å² SASA gate, we count cross-chain Cα neighbors within 10 Å, add small jitter (±1), cap at 12, and normalize to [0,1]. During training, a subset of hotspots are written to the numeric hotspot_values channel (instead of the boolean mask), so residues with the densest 10 Å cross-chain contacts receive values near 1 and are emphasized more strongly. This preserves the discrete hotspot definition while providing graded “super hotspot” strength to steer designs toward tightly packed interface cores.

To limit context while preserving local geometry, we radially cropped the target around a randomly selected binder hotspot, retaining residues within >25 Å of that hotspot (binder residues were always retained). Unstructured binder termini outside the interface were automatically trimmed using a sliding 9-residue window to measure side-chain neighbor support (sc_neigh threshold = 1.0). Terminal segments were removed if they contained ≤0/1/2 folded residues for windows of length ≤4/≤8/>8, respectively. If intervening residues between the cropped region and the hotspot bounds remained unfolded, trimming was permitted to extend slightly into the hotspot region. This procedure focused the model on compact, physically plausible binder-target interfaces while avoiding irrelevant or disordered regions. During training, the diffusion noise is centered at the origin (ORI), and the model learns correlation between the binder’s center of mass and the center of that noise cloud. At inference we exploit this by choosing the ORI location to steer where the binder is placed (specified by a pseudoatom in the inference inputs). Placing ORI at the geometric center of the hotspot mask yielded the most consistent interface placement, though positioning it near the anticipated binder site also produced robust, shape-complementary designs.

In total three models were trained using different strategies. One model was fine-tuning from RF2 structure prediction weights (ppi_robust_struct) and two from pre-trained RFD2 weights: RFD_45 (ptm_finetuned) and RFD_140 (ppi_robust). Training examples for ppi_robust and ppi_robust_struct were filtered for well folded structures as described above. ptm_fintuned was trained with additional examples of phosphorylation binding motifs from the pdb that were previously excluded during training with additional focus on masking target coordinates in some examples. This allows to generate both binder structure and target conformation at the same time, similar as in RFDiffusion (https://github.com/RosettaCommons/RFdiffusion/blob/main/examples/design_ppi_flexible_pepti <u>de.sh</u>) and Liu et al. (28).

### Binder design and prediction

Protein binders targeting a specific phosphopeptide sequence were computationally designed using a multi-stage protocol. Initial binder scaffold generation was performed with RFdiffusion (using the AA_ppi model where applicable), constraining the binder length to fewer than 160 amino acids. During this process, the target phosphopeptide sequence was held fixed, while its coordinates were allowed to co-diffuse with the *de novo* binder backbone. Following backbone generation, binder sequences were designed using LigandMPNN. For generated backbones that include a predefined structured motif (i.e., short peptide-protein interface coordinate) generated from Logos on a 4-7 amino acid peptide side, the key motif residues were held fixed during this sequence design step. The resulting protein-peptide complex structures were refined using the Rosetta FastRelax protocol, followed by a subsequent round of LigandMPNN to optimize the sequence for the relaxed structure. At this stage, an initial filtering step was applied to remove any design that did not form at least two hydrogen bonds with the phosphate group of the target phosphotyrosine residue. Finally, the structures of promising binder-peptide complexes were predicted using AlphaFold3 resulting in the final design model.

### Iterative sequence diversification

To expand the pool of potential candidates, AlphaFold3 predictions that passed the filtering criteria (detailed below) were subjected to an additional round of sequence design. The predicted backbone conformations of the binder-peptide pairs were held rigid, and LigandMPNN was utilized to generate sequence variants for the binder scaffold. This iterative technique aimed to increase sequence diversity for promising backbones. The newly generated sequences were then re-submitted for AlphaFold3 structure prediction and subsequently evaluated against the established filtering metrics.

### Design filtering criteria

Predicted binder-peptide structures from AlphaFold3 were rigorously filtered based on a combination of model confidence metrics and specific structural features. Designs were retained only if they met the following criteria:

1. Model Confidence: High per-residue confidence for the binder (high pLDDT), low inter-domain predicted aligned error between the binder and peptide (low iPAE), high inter-domain predicted TM-score (high iPTM), and a high overall predicted TM-score for the binder (high PTM).
2. Binding Specificity: To ensure focused interaction with the phosphotyrosine, designs were required to have at least two hydrogen bonds to the phosphotyrosine’s phosphate group. To penalize non-specific or excessive non-phosphorylated binding, designs with more than twelve total hydrogen bonds to the peptide were excluded.
3. Motif Fidelity: For designs built around a structural motif, the root-mean-square deviation (RMSD) of the phosphotyrosine residue between the designed and predicted structures was required to be less than 5 Å. This criterion ensures the target residue remains correctly oriented within the intended binding pocket.

In our experience, successful designs had a low iPAE between the binder and the phospho-residue of the peptide, and a high pLDDT for the target phospho-peptide, which is indicative of a highly confident prediction for the binding mode. In filtering sequence-backbone pairs we also took inspiration from the fact that several known phospho-binding motifs form extensive hydrogen bond networks to the phosphate group (e.g. SH2 domains), and as expected the crystal structures match the AF3 prediction of the designed sequence (13, 29).

### DNA library preparation

Sequences were padded with a (GGGS)n linker at the C terminal of designs to avoid biased amplification of short PCR fragments during PCR. Selected binder protein sequences were reverse translated using DNAworks2.0 for optimal expression in *S.cerevisae.* Oligo pools for the *de novo* binders were obtained from Twist Bioscience. Pools were amplified with Kapa HiFI Polymerase (Kapa Biosystems) using a qPCR machine (BioRAD CFX96). First, pools were amplified in a 25 uL reaction and terminated at half the maximum yield, avoiding over-amplification. PCR product was then loaded onto a DNA agarose gel, excised, and extracted with QIAquick Clean Up Kit (Qiagen, Inc.). 2-3 micrograms of pETCON vector and 6 micrograms of amplified DNA was transformed into yeast.

### Peptide synthesis

Protected phospho amino acids (e.g. Fmoc-Tyr(PO(OBzl)OH)-OH) were purchased from Millepore Sigma. Phosphopeptides were synthesized on a 0.1mmol scale via microwave-assisted solid-phase peptide synthesis (SPPS) using a CEM LibertyBlue peptide synthesizer and standard SPPS protocols. Peptides were subsequently coupled with Fmoc-AHX-OH (P3 Biosystems, 3eq.), HATU (Millepore Sigma, 3eq.), and N,N-dimethylformamide (DIEA, Millepore Sigma, 5eq) in DMF for 3 hours, then treated with 20% piperidine (Millepore Sigma) in DMF for 2 x 15 minutes, washed thoroughly with DMF and swelled with 50/50 DMSO/DMF for 30m prior to coupling with D-Biotin (Millepore Sigma, 3eq), HATU (3eq), DIEA (5eq) in 50/50 DMSO/DMF for 3 hours. The labeled peptide was then globally deprotected with a cleavage cocktail of 92.5/2.5/2.5/2.5 TFA/TIPS/H2O/2,2-(ethhylenedioxy)diethanetihol for 3h, concentrated briefly *in vacuo*, and precipitated into ice cold diethyl ether. The crude peptide was washed with ether and dried under nitrogen, then was resuspended in a minimal amount of water and acetonitrile (ACN) and purified via RP-HPLC (A: H2O with 0.1% TFA, B: ACN with 0.1% TFA) in a linear gradient from 5%B to 45%B. Fractions were collected and analyzed for product using an Agilent G6230B TOF, then pooled and lyophilized to yield pure phosphopeptide.

### Yeast surface display

A yeast surface display library was constructed using S. cerevisiae strain EBY100 transformed with a plasmid pool encoding the binder variants. Initial cultures were grown overnight at 30 °C in a selective synthetic complete medium lacking tryptophan and uracil (C-Trp-Ura), supplemented with 2% (w/v) glucose. For protein expression, the culture was diluted 1:10 into SGCAA auto-induction medium containing 0.2% (w/v) glucose and incubated overnight at 30 °C. Following induction, cells were washed with chilled PBS supplemented with 1% (w/v) Bovine Serum Albumin (PBSF). To confirm and enrich for clones expressing the binder construct, an initial sort was performed. Cells were labeled with a fluorescein isothiocyanate (FITC)-conjugated anti-c-Myc antibody (Miltenyi Biotec), which targets a c-Myc epitope tag on the displayed protein. The FITC-positive population was collected via FACS and regrown to generate an enriched expression library for subsequent affinity-based sorting.

Cells were regrown and washed, then labelled with biotinylated target (peptide) using two techniques - with avidity and without avidity. With avidity labeling, cells were incubated with biotinylated target, anti-c-Myc fluoresceinisothiocyanate (FITC, Miltenyi Biotech), and streptavidin-phycoerythrin (SAPE, ThermoFisher). Concentration of SAPE with the avidity method was 1/4 of the biotinylated target concentration. After the first few rounds of screening using the with-avidity method, weak binder candidates were eliminated. Using the without-avidity method, cells were incubated with a biotinylated target, washed, and then incubated with SAPE and FITC. Subsequent sorts were applied with varying concentrations.

### Competition-based sorting for phosphopeptide specificity

In some instances, to isolate binders with high specificity for the phosphorylated peptide, a competitive sorting strategy was employed. The induced and washed yeast library was first incubated with 300 nM of a biotinylated, unphosphorylated “decoy” peptide. Following a wash step, free biotin was added to the cell suspension to quench any unbound streptavidin-conjugated reagents and minimize non-specific interactions in the subsequent step. Next, cells were incubated with 100 nM of the biotinylated phosphopeptide “target.” To detect binding, the unphosphorylated peptide was labeled with an anti-biotin antibody conjugated to Allophycocyanin (APC), while the phosphopeptide was labeled with an anti-biotin antibody conjugated to Streptavidin-Phycoerythrin (SAPE). Using FACS, distinct cell populations were gated and sorted;

1. Specific Binders: High SAPE signal and low/negative APC signal.
2. Non-specific Binders: High APC signal.
3. Dual-Recognition Binders: High signal in both SAPE and APC channels.

Populations showing specific binding to the phosphopeptide were collected, regrown, and subjected to up to three additional rounds of sorting to achieve high enrichment.

### Isolation and Sanger sequencing of individual clones

The enriched cell population from the final sorting round was plated on solid agar medium for yeast growth and incubated overnight at 30°C. Individual colonies were isolated, and the plasmid DNA was prepared for amplification via colony PCR. A small portion of each colony was boiled to lyse the cells, and the binder-encoding DNA insert was amplified using vector-specific primers (pETCON primers). PCR products of the expected size were purified and analyzed by Sanger sequencing to determine the DNA sequence of individual high-performing protein binders.

### Next-generation sequencing (NGS) of enriched populations

To comprehensively analyze the convergence of the selected binder pool, an aliquot of the sorted cells from each round of enrichment was collected for NGS analysis. Plasmid DNA was extracted from the bulk sorted populations using a commercial extraction kit (Zymoprep, Zymo Research). The variable binder-encoding regions were then amplified from the extracted plasmid pool via PCR using primers that both flanked the insert and appended Illumina-compatible sequencing adapters and indices. The resulting amplicon libraries were purified, quantified, and sequenced on an Illumina platform (NextSeq/MiSeq). This deep sequencing approach allowed for the quantitative tracking of sequence diversity and the enrichment of specific binder families throughout the selection campaign.

### Bacterial protein production and purification

*E. coli* Lemo21(DE3) strains were transformed with the pET29b+ plasmid encoding the binder gene. Cells were then grown for 24 hours in liquid broth medium supplemented with kanamycin and inoculated at 1:50 mL ratio in the Studier TBM-5052 autoinduction medium supplemented with kanamycin for several hours at 37°C, then grown overnight at 18°C. Cells were collected by centrifugation at 4,000 g at 4°C for 15 minutes, resuspended in 30 mL of lysis buffer (20 mM Tris-HCl, pH 8.0, 300 mM NaCl, 30 mM imidazole, 1 mM PMSF and 0.02 mg ml-1 DNase). The cell resuspension was lysed by sonication for 2.5 minutes (5 second cycles) and the lysate was clarified by centrifugation at 24,000 g at 4°C for 20 minutes. The supernatant was passed through a Ni-NTA nickel resin, pre-equilibriated with a wash buffer (20 mM Tris-HCl, pH 8.0, 300 mM NaCl and 30 mM imidazole). The resin was washed twice with 10 column volumes of wash buffer and eluted with 3 column volumes of elution buffer (wash buffer with 300 mM imidazole). Eluted proteins were concentrated using Ultra-15 Centrifugal Filter Units and purified with Superdex 75 Increase 10/300 GL size exclusion column in TBS (25 mM Tris-HCl, pH 8.0, and 150 mM NaCl). Fractions containing monomeric proteins were pooled, concentrated and snap-frozen.

### Direct production of proteins from designs (30)

Linear gene fragments encoding the binder design sequences were cloned into the LM627 vector using Golden Gate assembly. These subcloning reactions were performed in 96-well PCR plates with a 4 µL reaction volume. Following this, 1 µL of the reaction mixture was transformed into chemically competent E. coli BL21 (DE3) cells. After a 1-hour recovery period in 100 µL of SOC medium, the transformed cell suspensions were transferred directly into a 96-deep well plate containing 900 µL of LB media supplemented with Kanamycin. After overnight incubation at 37°C, 100 µL of the growth culture was inoculated into 96-deep well plates containing 900 µL of auto-induction media (autoclaved TBII media with Kanamycin, 2 mM MgSO4, 1X 5052). The cultures were then expressed overnight at room temperature. Cells were harvested by centrifugation at 4000 x g for 15 minutes. The bacterial pellets were lysed in 100 µL of lysis buffer (1X BugBuster (Millipore #70921-4), 0.01 mg/mL DNase, 1 Pierce protease inhibitor tablet per 50 mL lysis) for 30 minutes on shaker, 220 RPM. The lysates were spun down by centrifugation at 4000 x g for 10 minutes, followed by purification using Ni-charged MagBeads (GenScript #L00295). The wash buffer contained 25 mM Tris pH 8.0, 300 mM NaCl, and 10 mM Imidazole, while the elution buffer contained 25 mM Tris pH 8.0, 300 mM NaCl, and 500 mM Imidazole. 250 µL of wash buffer was used to wash twice while 120 µL of elution buffer was used to collect eluted proteins. Filtered elutions were then submitted to HPLC, S200. For samples showing major monomeric peaks, protein concentrations were determined by measuring absorbance at 280 nm with a NanoDrop spectrophotometer (Thermo Scientific), using extinction coefficients and molecular weights calculated from their amino acid sequences.

### Biolayer interferometry

Protein-peptide interactions were measured using an Octet RED96 System (ForteBio) with streptavidin-coated biosensors (ForteBio). Wells containing 200 uL of solution, and HBS-EP+ buffer (GE Healthcare Life Sciences, 10 mM HEPES pH 7.4, 150 mM NaCl, 3 mM EDTA, 0.05% (v/v) surfactant P20) plus 1% non-fat dry milk blotting grade blocker (BioRad). Biosensor tips were loaded with target peptide at 5µM for 300 s (threshold of 0.6 nm response), then incubated in the HBS-EP+ buffer for 60s for baseline measurement. Tips were then dipped into the solution containing binder for 600s (association) and finally, dipped into the HSP-EP+ buffer for 600s (dissociation). Binding data was analyzed with ForteBio Data Analysis Software version 9.0.0.10.

### Protein crystallography and structure determination

Purified protein was used for crystallization trials using the sitting drop vapor diffusion method. Crystallization trials were set up by Mosquito LCP from SPT Labtech. Formulated total 200 nL drops using the 96-well plate format at 20°C. Crystal search was conducted on plates using UV imaging (UVEX microscopes and UVEX PS-256 from JAN Scientific). Diffraction quality crystals were obtained with the following conditions;

1. CD3ε-pY188 crystals grew in 2.0 M Sodium chloride and 10% (w/v) PEG 6000.
2. EGFR-pY1068 crystals grew in 0.1 M Na HEPES pH 7.0 and 2 M ammonium sulfate

Diffraction data was obtained either at the National Synchrotron Light Source II on beamline 17_ID-1 (AMX) or Advanced Photon Source on beamline NECAT 24ID-E. XDS was used to integrate x-ray intensities and subsequent data reduction (31). Data was merged and scaled using Pointless/Aimless in the CCP4 program suite (32). Molecular replacement (Phaser program) was used for structure determination and phasing (33). Models were improved using Phenix autobuild (rebuild-in-place set to false, using simulated annealing (34)). Manual model building was performed using COOT (35). The final model was evaluated using MolProbity (36).

### Reagents

The JaneliaFlour dye JFX-554 was a gift from Luke Lavis. Carrier free EGF was purchased from R&D Systems (#236-EG) and was reconstituted following the manufacturer’s protocol to a concentration of 200 µg/mL. Afatinib was purchased from SelleckChem (#S1011), dissolved in DMSO (ThermoFisher, #D12345) to a concentration of 10 mM, and diluted directly into cell culture medium for incubation at a final concentration of 10 µM.

### Tissue culture and transfections

CHO cells were electroporated using the Lonza SF Cell Line 4D-Nucleofector X Kit L (Lonza, # V4XC-2024) following the CHO-K1 protocol (1 million cells, 2 μg of DNA) in a 4D-Nucleofector (Lonza). Following electroporation cells were moved into a microcentrifuge tube containing ∼0.9 mL of RPMI (Gibco, #A10491-01) with 10% FBS (Genesee Scientific, 25-514H) and incubated for 10 minutes at 37°C. Following incubation, approximately 250,000 cells from the suspension were transferred to 35 mm poly-D-lysine coated FluoroDish optical dishes (WPI, #FD35PDL-100) with pre-warmed complete media for final volume of ∼2 mL. Prior to cell seeding, optical dishes were cleaned using 1 M KOH (Fisher Chemical, #AA46072AP) for 30 minutes, rinsed with Milli-Q H_2_O 5 times, incubated with 100% ethanol (Decon Labs, #2705HC) for 30 minutes, followed by another 5 rinses with Milli-Q H_2_O.

### Plasmids and cloning

All constructs were cloned using Platinum SuperFi II PCR Master Mix (Invitrogen, #12368010) following manufacturer’s protocols. EGFR-GFP was purchased from AddGene (AddGene plasmid #32751), Halo-EGFR was generated as previously described (37). The EGFR^Y1068S^ construct was made using the EGFR-GFP plasmid and overlapping primers containing the single base-pair mutation and ∼15 base pairs of overlap (see table below). The resultant PCR product was incubated with KLD Enzyme Mix (NEB, #M0554S) at 25°C for 5 minutes,transformed into chemically competent DH5α *E. coli* cells (ThermoFisher, #FEREC0112) and plated onto LB agar plates containing 50 µg/mL of kanamycin (ThermoFisher, #J61272.09). Multiple colonies were screened by whole plasmid sequencing to identify constructs containing the desired mutation.

pCDNA5/FRT/TO_H2B_HaloTag7 was purchased from AddGene (AddGene, plasmid #169329) and the subsequent EGFR_pY1068_bp construct was made using the Gibson assembly approach (see table for primers) to add HaloTag7 to the C-terminus of binder pY1068. After PCR amplification the products were combined using Gibson Assembly HiFi Master Mix (ThermoFisher, #A46627). PCR product was then transformed into chemically competent DH5α *E. coli* cells (ThermoFisher, #EC0112) and plated onto LB agar plates containing ampicillin (ThermoFisher, #J63807-09). Multiple colonies were screened by whole plasmid sequencing to identify plasmids containing the desired construct.

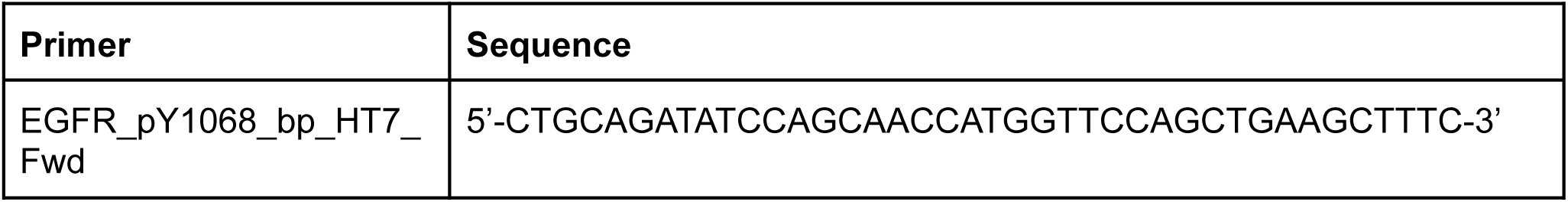
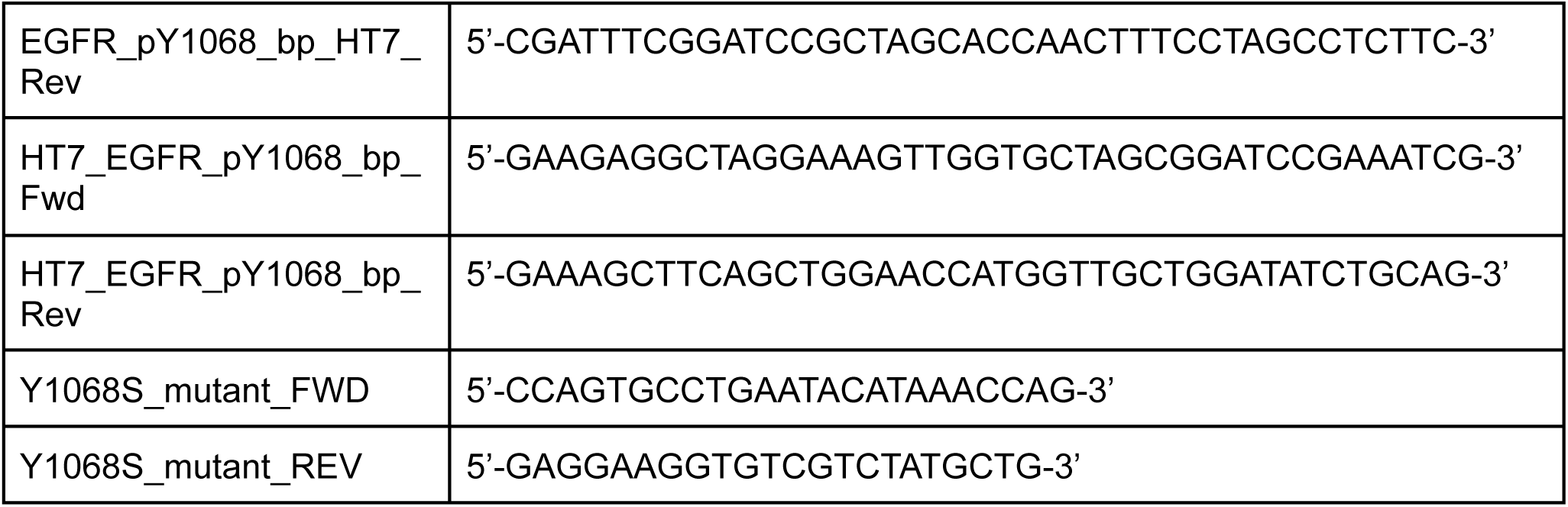

### Single-particle tracking (SPT)

Single-particle tracking was carried out using a Nikon Ti-2 STORM/TIRF inverted microscope equipped with a 100x 1.49 NA oil immersion objective with an objective heater and Tokai Hit stage-top incubator, both kept at ∼37°C. A 561 nm laser (17.77 W/cm^2^ power at the optical dish) was used for excitation of JFX-554 using a ZT 405/488/561/640 nm (Chroma, #88901) filter cube. Emission was detected using two Andor iXon Life 897 EMCCDs (pixel size 0.160 μm at 30.85Hz), and emission was split to each camera using a TwinCam (Cairn), a 640 nm LP dichroic filter, with a 600/40 nm BP filter, and a 655 nm LP filter.

Electroporated CHO cells were plated the day before imaging in 35 mm FluoroDishes using complete medium. For experiments using afatinib, cells were incubated with the TKI at a concentration of 10 μM for 1 hr at 37°C in complete media. For all imaging experiments cells were incubated with 50 nM JFX-554 for 10 minutes at 37°C. Following labeling, excess dye was washed out using PBS (Gibco, 14040-133). Cells were then imaged in ‘Live Cell Imaging Solution’ (Invitrogen, #A14291DJ) supplemented with 10 mM glucose (ThermoFisher, #D16-500) at 37°C. To ensure co-transfection prior to imaging cells in plates transfected with eitherEGFR-GFP or EGFR^Y1068S^ were briefly checked for GFP expression using a 488 nm laser (16.41 W/cm^2^ power at the optical dish) for excitation. For all experiments 3 cells in each dish were first imaged for 5 minutes without ligand. Then ligand was added to the live cell imaging buffer at a final concentration of 50 nM and allowed to incubate for 5 minutes, then 3 cells were imaged for the +EGF conditions. The first minute of all recorded videos was used for photobleaching and the following 4 minutes were used for data analysis.

### EGFR_pY1068_bp^HT7^ Tracking

EGFR_pY1068_bp^HT7^ was tracked using TrackIt in MATLAB, a software for analyzing single-molecule fluorescence microscopy data. TrackIt is available from the Gebhardt Lab (https://gitlab.com/GebhardtLab/TrackIt) (38). Movies in TIFF format were loaded into the TrackIt and regions of interest (ROIs) were drawn around cells. Parameters for tracking were set as the following: threshold factor of 2, tracking radius (px) of 4.3, minimum track length of 20, gap frames of 10, and a minimum track length before gap frame of 2. Following tracking in TrackIt Divide-and-Conquer Moment Scaling Spectrum transient diffusion analysis (DC-MSS) software from Khuloud Jaqaman’s lab (https://github.com/kjaqaman/DC-MSS) was used to distinguish between diffusion classes of tracked binders including immobile, confined, and free tracks (17).

Data outputs from DC-MSS analysis were graphed and analyzed using Origin 2025b (10.2.5.234). MATLAB version 2025b (25.2.0.3042426) was used to run TrackIt and DC-MSS.

## Supplementary Note

### Detailed results of initial testing of designs

We obtained synthetic DNA encoding 459 designs for CD3ε pY188 (CD3ε_pY188_bp), 5,082 designs for EGFR pY1068 (EGFR_pY1068_bp), 2,408 designs for EGFR pY1173 (EGFR_pY1173_bp), and 80 designs for INSR pY1361 (INSR_pY1361_bp; the latter generated with the Logos pipeline). For the CD3ε and EGFR binders, we displayed the designed proteins on yeast, and evaluated binding to both pY and unmodified targets. To do this, we incubated the yeast libraries with biotinylated non-phosphorylated peptides and streptavidin-Allophycocyanin (APC, far-red color), washed several times, incubated with excess biotin to block unbound Streptavidin-APC, and then incubated with biotinylated phosphorylated peptides and streptavidin-phycoerythrin (PE, red color). Designs specifically binding the phosphorylated target after several rounds of sorting were identified by sequencing. For the INSR binders, we obtained the individual genes encoding the designs and identified binders through direct *in vitro* purification and binding assays (see Methods for details).

For CD3ε, phosphorylation-dependent binders were identified by Sanger sequencing of 47 single clones grown from competition sort three (see Methods for more information). The sequences were closely related and one was cloned into plasmid pET29b+ for expression in E. coli. The CD3ε Y188 binder was assessed using biolayer interferometry (BLI) and found to have a Kd of 2.3 μM to the phosphorylated CD3ε_pY188_bp peptide target and undetectable binding for the non-phosphorylated version of the peptide (**Figure 2A-C, row 1**).

## Author Contributions

Conceptualization and overall study design: A.L., A.M., B.C., D.B., D.T., F.D., G.R.L., H.A.T., J.Z.Z., K.W., M.S.B., P.S., R.K., W.A.; Contribution of code and ideas: B.C., D.J., D.T., H.A.T., M.S.B., P.S., R.I.B., R.K., W.A., J.Y.; Model training and optimization: B.C., D.T., G.R.L., H.A.T., J.Y., M.S.B., W.A.; Pipeline development for phosphopeptide binder design: G.R.L., I.K., J.Z.Z., K.A.K., K.W., M.S.B.; Computational binder design and structural analysis: G.R.L., J.S., J.Z.Z., K.A.K., K.W., L.C.H., M.S.B., R.K.S., S.Z.; Peptide synthesis and reagent preparation: C.M., X.L.; Yeast library construction: D.K.V., N.R.; Yeast display screening and selection: G.R.L., J.S., J.Z.Z., K.A.K., M.S.B., R.K.S., S.Z.; Protein expression and purification: G.R.L., J.Z.Z., K.A.K., K.W., M.S.B.; Biophysical characterization: G.R.L., J.S., J.Z.Z., K.A.K., M.S.B.; Crystallization, data collection, and structural determination: A.K., A.K.B., E.J.; Cloning and reagent preparation for live-cell imaging: I.M.S., K.C.M.; Single-particle tracking and data analysis: I.M.S., K.C.M.; Supervision and project administration: D.B., F.D., J.Z.Z., L.S., M.S.B.; Writing - original draft and review & editing: B.C., D.B., G.R.L., J.Z.Z., K.A.K., K.W., M.S.B., I.M.S., K.C.M., with input from all authors.

## Code and Data Availability

Code for RFD2-MI model inference and the phosphotyrosine binder design workflow is available at https://github.com/RosettaCommons/RFDiffusion2_all_the_code. Additional data and materials will be made available upon publication in accordance with journal policies.

## Acknowledgements

We thank L. Goldschmidt and K. VanWormer and their teams for maintaining the computational and wet lab resources at the Institute for Protein Design.

We gratefully acknowledge support from the Bill and Melinda Gates Foundation (INV-043758), the Defense Threat Reduction Agency (HDTRA1-19-1-0003, HDTRA1-21-1-0038), and a gift from Microsoft. This work was further supported by the Helmsley Charitable Trust Type 1 Diabetes Program (2019PG-T1D026), the Howard Hughes Medical Institute, and the Human Frontiers Science Program (RGP0061/2019).

We thank Spark Therapeutics for support through the project *Computational Design of a Half Size Functional ABCA4*. We also acknowledge the Audacious Project at the Institute for Protein Design, the Open Philanthropy Project Improving Protein Design Fund, and the Open Philanthropy Project Universal Flu Vaccine Fund.

This material is based upon work supported by the National Science Foundation (CHE-2226466). Additional support came from the National Institutes of Health, including the National Cancer Institute (R01CA240339, K99 CA293001, R00 CA256350), and the National Institute on Aging (R01AG063845).

This work was delivered in part as part of the MATCHMAKERS team supported by the Cancer Grand Challenges partnership funded by Cancer Research UK (CGCATF-2023/100008), the National Cancer Institute (OT2CA297288), and the Mark Foundation for Cancer Research. We also acknowledge support from the Advanced Research Projects Agency for Health (APECx Program, 1AY1AX000036), and the Washington Research Foundation Innovation Fellows Program.

This research used beamline 17_ID-1 (AMX) of the National Synchrotron Light Source II, a U.S. Department of Energy (DOE) Office of Science User Facility operated for the DOE Office of Science by Brookhaven National Laboratory under Contract No. DE-SC0012704. The Center for BioMolecular Structure (CBMS) is primarily supported by the National Institutes of Health, National Institute of General Medical Sciences (NIGMS) through a Center Core P30 Grant (P30GM133893), and by the DOE Office of Biological and Environmental Research (KP1605010). This work is based upon research conducted at the Northeastern Collaborative Access Team beamlines, which are funded by the National Institute of General Medical Sciences from the National Institutes of Health (P30 GM124165). The Eiger 16M detector on the 24-ID-E beam line is funded by a NIH-ORIP HEI grant (S10OD021527). This research used beamtime awards (DOI: https://doi.org/10.46936/APS-189869/60014006) from the Advanced Photon Source, a U.S. Department of Energy (DOE) Office of Science User Facility operated for the DOE Office of Science by Argonne National Laboratory under Contract No. DE-AC02-06CH11357

