## Supplementary Materials for "*De novo* design of phosphotyrosine peptide binders"

**Figure S1:**

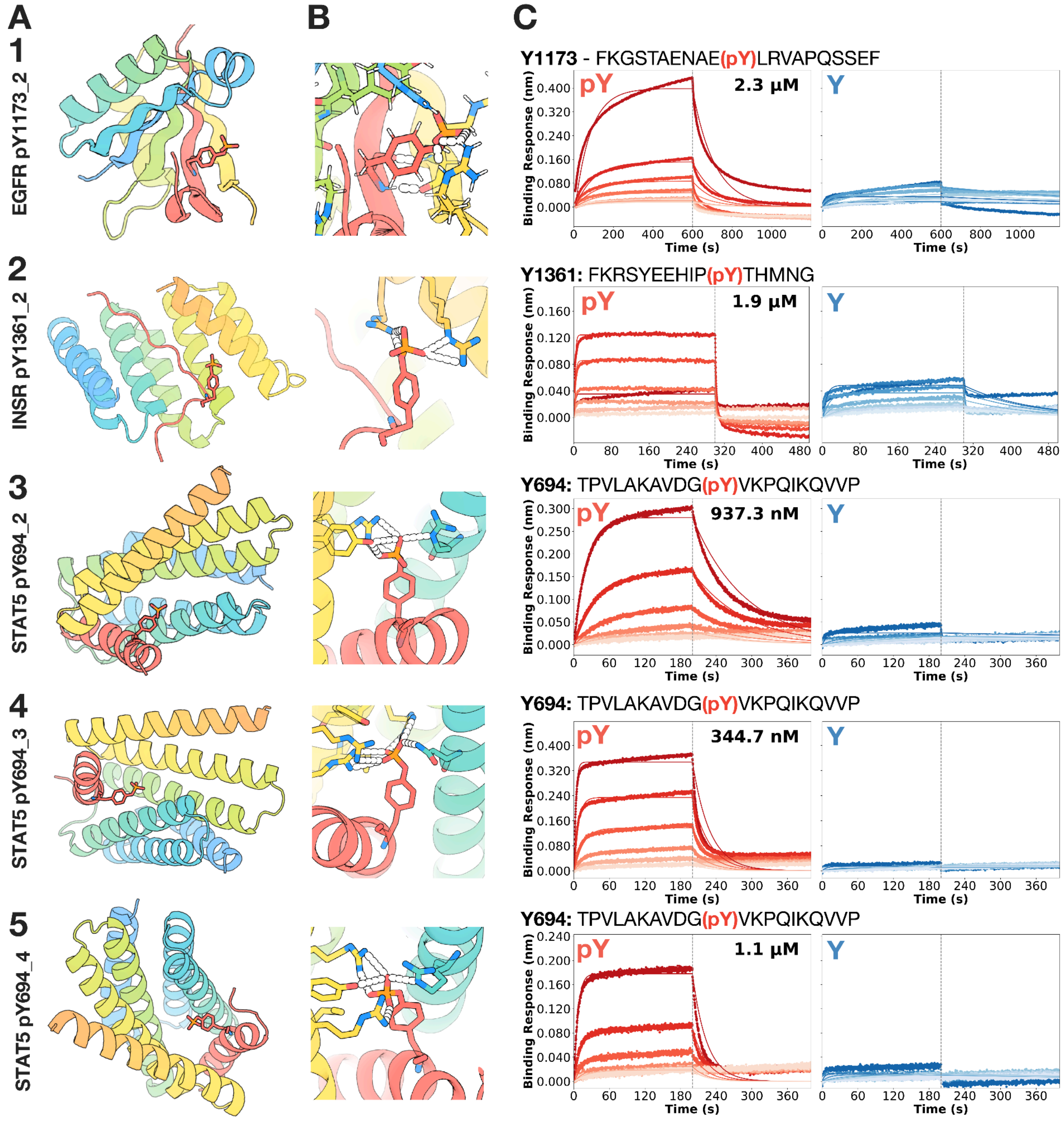


**Figure S1: Additional phosphopeptide binders.**

**(A)** Design models (rainbow) for the following target peptides: EGFR pY1173 (row 1), INSR (row 2) and STAT5 (row 3-5). In each case the phosphorylated peptide (red/orange sticks) docks into the designed pocket with high correspondence between observed and predicted backbone and side-chain geometry. **(B)** Close-up views of key hydrogen-bonding networks (dashed lines) in the predicted complexes, highlighting how the phosphotyrosine moiety (red/orange sticks) is anchored by a conserved polar pocket in the binder (rainbow). **(C)** Representative biolayer interferometry (BLI) sensorgrams showing association and dissociation of each binder to phosphorylated (blue) versus non-phosphorylated (red) peptides, with global 1:1 kinetic fits overlaid in black. They exhibit dose-dependent binding only to the phosphorylated peptide, showing a much more obvious binding toward pY peptides than their unmodified counterparts.

**Figure S2:**

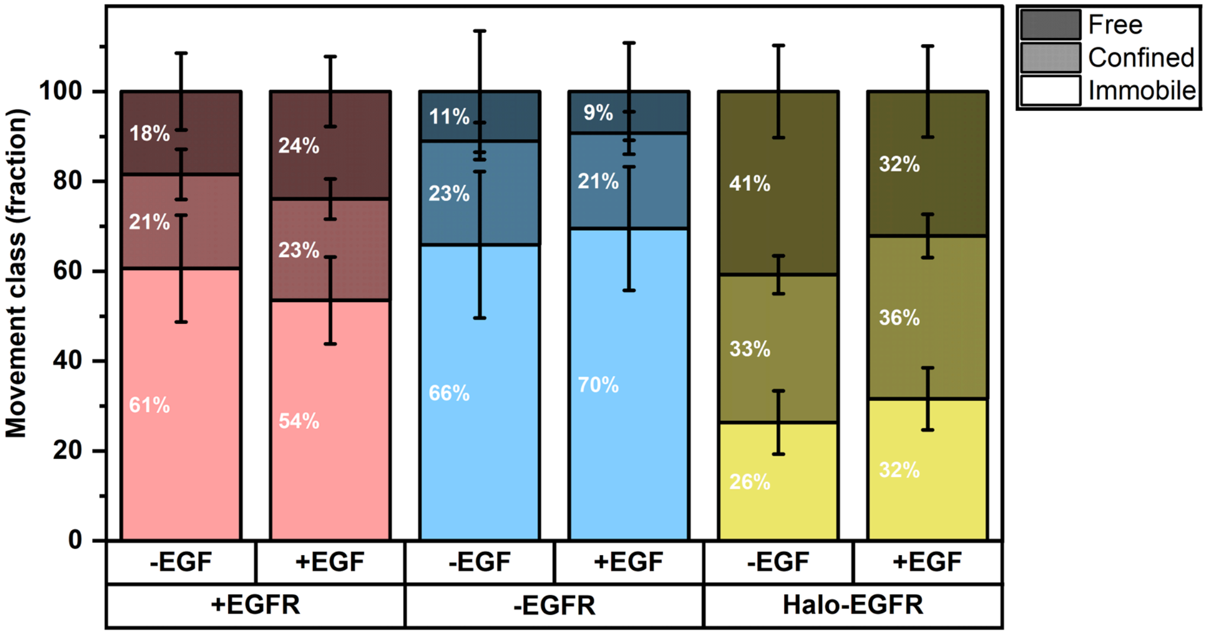


**Figure S2. EGFR pY1068 exhibits mostly immobile diffusion.** DC-MSS analysis was used to determine individual movement class fractions of EGFR_pY1068_bp$^{HT7}$ in cells expressing EGFR (pink), cells not expressing EGFR (blue), or cells expressing EGFR tagged with a HaloTag (yellow). Immobile tracks are shown with no shading, confined tracks are shown with medium shading, and free tracks are shown with dark shading. Error bars represent the standard deviation of the data.

**Figure S3:**

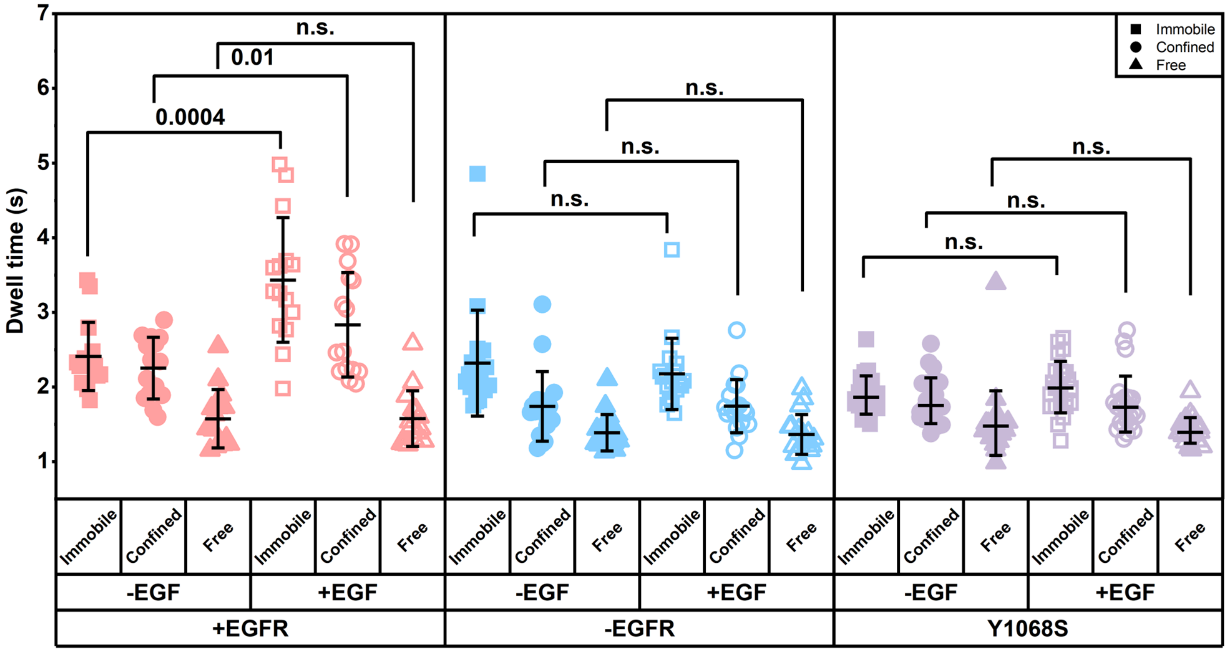


**Figure S3. Phospho-binder pY1068 shows increased dwell time at the plasma membrane when EGFR is stimulated with EGF.** DC-MSS analysis was used to parse EGFR_pY1068_bp$^{HT7}$ diffusion into immobile (square), confined (circle), and free (triangle) behavior. All CHO cells expressing EGFR_pY1068_bp$^{HT7}$, were co-transfected with wild-type EGFR (pink), not co-transfected with EGFR (blue), or were co-transfected with EGFR$^{Y1068S}$. Cells were either unstimulated (open symbols) or stimulated with 50 nM EGF (closed symbols). Unpaired two-sided Welche's t-tests were used to calculate p-values.
